# Distinct activities of Scrib module proteins organize epithelial polarity

**DOI:** 10.1101/866863

**Authors:** Mark J. Khoury, David Bilder

**Affiliations:** Department of Molecular and Cell Biology, University of California-Berkeley, Berkeley CA, 94720, USA

**Keywords:** Epithelia, polarity, Drosophila, Par complex, Scribble module

## Abstract

A polarized architecture is central to both epithelial structure and function. In many cells, polarity involves mutual antagonism between the Par complex and the Scrib module. While molecular mechanisms underlying Par-mediated apical determination are well-understood, how Scrib module proteins specify the basolateral domain remains unknown. Here, we demonstrate dependent and independent activities of Scrib, Dlg and Lgl using the *Drosophila* follicle epithelium. Our data support a linear hierarchy for localization, but rule out previously proposed protein-protein interactions as essential for polarization. Membrane recruitment of Scrib does not require palmitoylation or polar phospholipid binding but instead an independent cortically-stabilizing activity of Dlg. Scrib and Dlg do not directly antagonize aPKC, but may instead restrict aPKC localization by enabling the aPKC-inhibiting activity of Lgl. Importantly, while Scrib, Dlg and Lgl are each required, all three together are not sufficient to antagonize the Par complex. Our data demonstrate previously unappreciated diversity of function within the Scrib module and begin to define the elusive molecular functions of Scrib and Dlg.

**SIGNIFICANCE STATEMENT:** To enable their physiological functions, cells must polarize their plasma membrane. In many epithelia, polarity is regulated by balanced activity of the apical Par complex and basolateral Scribble module. While the former is understood in molecular detail, little is known about how the latter works. We identify distinct functions of the three Scribble module proteins, separating independent roles in a localization hierarchy from cooperative roles in apical polarity antagonism and showing that they are not together sufficient to specify basolateral identity. This work establishes an essential basis for a mechanistic understanding of this core polarity machinery that controls processes ranging from stem cell divisions to organ morphogenesis across animal species.

## INTRODUCTION

Cell polarity is defined by the coexistence of two distinct spatial identities within the confines of a single plasma membrane. This process is critical for many cell types, including stem cells, epithelial cells, migratory cells, and immune cells, to carry out their physiological functions (1, 2). Despite the distinct manifestations of polarity in these specialized cells, polarity in each is generated by a common pathway involving a set of conserved protein modules (3–5). Foremost among these are the Par and Scrib modules, consisting of Par-3, Par-6 and atypical Protein Kinase C (aPKC) for the former and Scribble (Scrib), Discs-large (Dlg) and Lethal giant larvae (Lgl) for the latter (3, 4). These proteins play crucial roles in diverse biological processes and have also been implicated in numerous pathologies, from congenital birth defects to cancer (3, 4, 6). Thus, uncovering their molecular activities is essential to a mechanistic understanding of cell, developmental and disease biology.

A number of studies have provided important insight into the molecular function of the Par module and each of its individual components (7–11). Much of this work derives from *Drosophila* epithelial cells and neural stem cells, where the Par module regulates the apical domain and the Scrib module is required to specify the basolateral domain. The core distinction of membrane domains arises from mutual antagonism between the two modules, centering around interactions between aPKC and Lgl (**Fig. 1A**). In the apical domain, aPKC phosphorylates Lgl on three residues within a polybasic domain, causing it to dissociate from the plasma membrane (12–14). Conversely, Lgl inhibits aPKC kinase activity and localization along the basolateral cortex (15–17). Many details of Par protein activities and their outcomes are now understood, including specific protein-protein interactions in dynamic complexes, their structural basis, post-translational modifications and the kinetic order of events during apical polarization (18, 19).

**Figure 1.**
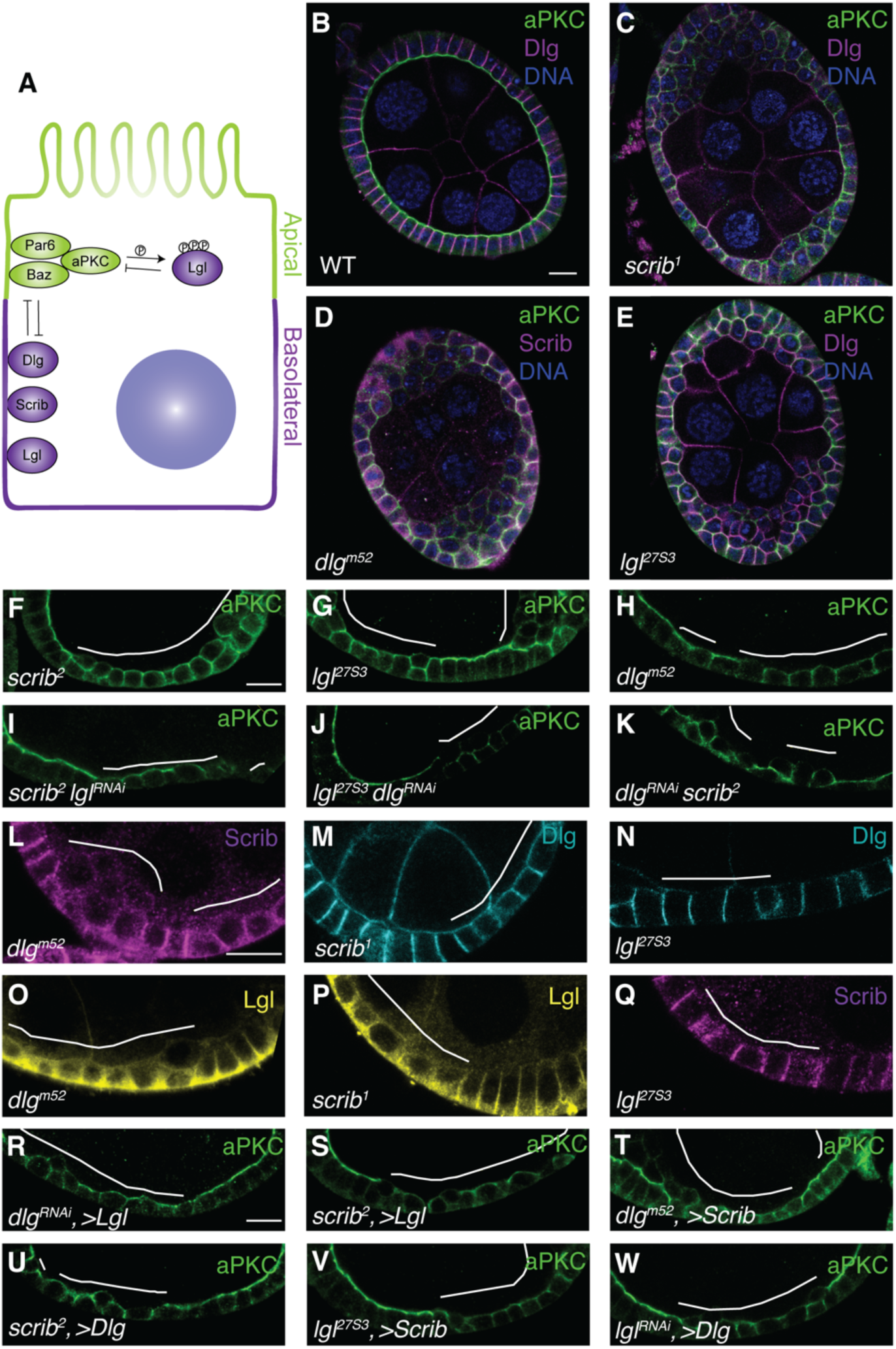
Functional relationships within the Scrib module. (A) Simplified schematic representation of epithelial polarity interactions. Polarity phenotypes of WT (B), *scrib* (C), *dlg* (D) and *lgl* (E) follicle cells: mutants exhibit polarity loss, characterized by mixing of apical and basolateral domains and multilayering of the epithelium. Compared to single mutants (F-H), double depleted combinations (I-K) do not show an enhanced apical expansion phenotype. Localization of Scrib module proteins: both Scrib and Lgl show hazy, cytoplasmic mislocalization in *dlg* mutant cells (L,O). In *scrib* mutant cells, Dlg localization is normal (M), while Lgl is mislocalized (P). In *lgl* mutants, both Scrib and Dlg localizations are unchanged (N,Q). Overexpression of Lgl does not rescue apical polarity defects in *dlg* or *scrib* mutants (R,S). Scrib overexpression cannot rescue *dlg* mutants (T) nor can Dlg overexpression rescue *scrib* mutants (U). *lgl* mutants are not rescued by Scrib or Dlg overexpression (V,W). Scale bars, 10µm. White line indicates mutant cells and/or overexpression clones in this and all subsequent figures.

By contrast to the wealth of mechanistic information about the Par complex, and despite the discovery of the relevant genes decades ago, the molecular mechanisms of basolateral domain specification by the Scrib module are still unknown. All three genes encode large scaffolding proteins containing multiple protein-protein interaction domains and lack obvious catalytic activity (13, 20–22). Recent studies have identified novel interacting partners of Scrib module proteins, but few of these interactors have been implicated as regulators of cell polarity themselves (23, 24). Moreover, few studies have focused on the regulatory relationships within the Scrib module itself, and beyond the well-characterized aPKC-inhibiting function of Lgl, the fundamental molecular activities of Scrib and Dlg remain unknown. In this work, we identify distinct activities of Scrib, Dlg and Lgl that are required but not sufficient for basolateral polarization, shedding light on the mechanisms that restrict the Par complex to partition the epithelial cell membrane.

## RESULTS

### A linear hierarchy for localization but not function of basolateral polarity regulators

We used the conserved epithelial features of *Drosophila* ovarian follicle cells to study regulation of the basolateral membrane domain (25). Cells mutant for null alleles of *scrib, dlg* or *lgl* lose polarity, characterized by mixing of apical and basolateral domains and cells form multilayered masses at the poles of the egg chamber (**Fig. 1B-E**). Importantly, we focused our analysis on the central follicle epithelium, where polarity-deficient cells retain relatively normal morphology that allows accurate monitoring of protein localization. We first asked whether Scrib, Dlg and Lgl have independent as well as shared functions in epithelial polarity. We generated follicle cells simultaneously mutant for one of the genes and expressing a validated RNAi transgene targeting a second gene. However, we saw no differences between single mutant cells and cells double-depleted for *scrib dlg, scrib lgl* and *lgl dlg*, and in no case was the aPKC mislocalization seen in a single mutant enhanced (**Fig. 1F-K**). These phenotypes are consistent with Scrib, Dlg and Lgl regulating polarity through a single, common pathway.

We next defined regulatory relationships between Scrib module components. Previous work in several organs has documented mutual dependence for localization, but also significant differences in their interrelationships (26–28). In *dlg* mutant follicle cells, both Scrib and Lgl are mislocalized and exhibit hazy, cytoplasmic distributions (**Fig. 1L,O**). In *scrib* mutant follicle cells, although Lgl is mislocalized as in *dlg* mutants, Dlg maintains normal basolateral localization (**Fig. 1M,P**). Moreover, in *lgl* mutant follicle cells, both Scrib and Dlg maintain normally polarized cortical domains (**Fig. 1N,Q**). These results suggest a linear pathway whereby Dlg localizes independently to the plasma membrane, Scrib localization requires Dlg, and Lgl localization is dependent on both other Scrib module proteins.

We then asked whether elevated levels of one protein in this pathway could compensate for loss of another. We first tested overexpression of Lgl in *scrib* or *dlg* mutant cells and found that this did not modify the phenotype of either mutant (**Fig. 1R,S**). Similarly, Scrib overexpression did not modify the *dlg* mutant phenotype, and Dlg overexpression did not modify the *scrib* mutant phenotype (**Fig. 1T,U**). Moreover, neither Scrib nor Dlg overexpression was able to modify the *lgl* mutant phenotype (**Fig. 1V,W**). These data suggest that, despite the linear localization hierarchy, regulation of basolateral polarity involves relationships that cannot be bypassed by simple overexpression of one Scrib module component.

### Dlg stabilizes Scrib at the cortex

Since Dlg is required for Scrib cortical localization, we investigated the underlying mechanism. We used Fluorescence Recovery After Photobleaching (FRAP) assays to compare the stabilities of each protein at the plasma membrane, using functional GFP-tagged versions expressed from endogenous loci. In WT cells, Scrib::GFP was highly stable, whereas Dlg::GFP was intermediately dynamic and Lgl::GFP was comparatively mobile (**Fig. 2A**). Strikingly, in *dlg*-depleted cells, Scrib::GFP exhibited an approximately fourfold increase in recovery kinetics, consistent with the loss of cortical localization also seen in fixed tissue (**Fig. 2B**). By contrast, although Dlg::GFP in *scrib* and *lgl* mutant cells remained localized at the cortex and mobile fractions are not changed, it also exhibited increased recovery kinetics (**Fig. S1A,B**), perhaps reflecting increased in-plane mobility due to defective septate junction formation (29, 30). Importantly, however, Scrib::GFP was unchanged in *lgl* mutant cells (**Fig. S1C**). Thus, FRAP assays support an important role for Dlg in stabilizing Scrib on the plasma membrane.

**Figure 2.**
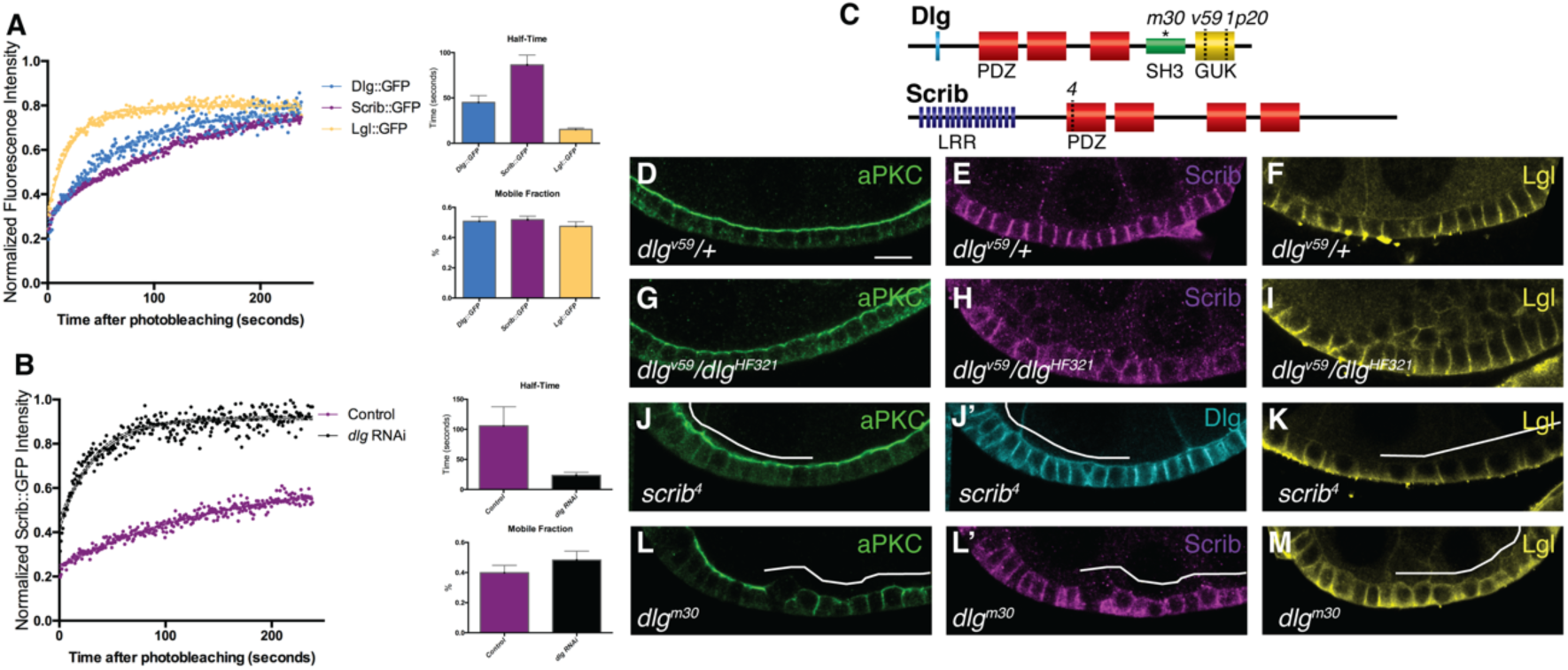
Dlg regulates cortical Scrib stability. FRAP assay (A) shows distinct mobility of Scrib, Dlg and Lgl in WT cells; Scrib is most stable and Lgl is most dynamic. (B) Scrib shows a ∼4 fold increase in recovery kinetics in *dlg* RNAi cells. (C) Schematic of the SH3 and GUK mutant *dlg* alleles and PDZ mutant *scrib* alleles used in D-M. Compared to WT (D-F), aPKC (G) and Lgl (I) localizations are unaffected in cells mutant for a *dlg* GUK-deficient allele. Scrib localization (H) is partially affected, although this may be due to reduced stability of mutant Dlg. aPKC (J), Dlg (J’) and Lgl (K) localizations are unaffected in cells mutant for a *scrib* allele lacking PDZ domains. (L) aPKC mislocalizes laterally in cells mutant for a *dlg* SH3 point mutant allele. Scrib (L’) and Lgl (M) also exhibit cytoplasmic mislocalization in these cells. Scale bars 10µm. Error bars represent 95% confidence intervals.

One mechanism that could localize Scrib to the cortex is a phospholipid-binding polybasic motif (PBM), as seen in other polarized proteins, including Lgl and aPKC (13, 14, 31). However, an obvious PBM is not seen in the Scrib protein sequence. PBMs directly bind polar phospholipids, but mutating PI4KIIIα or expressing dominant negative PI3K (Δp60), which deplete PIP2 and PIP3, respectively, did not alter Scrib plasma membrane localization (**Fig. S2A,B**)(32, 33). Additionally, ATP depletion by Antimycin A treatment, which reduces PIP levels and is sufficient to delocalize Lgl::GFP, did not alter Scrib::GFP localization (**Fig. S2C-F**)(13).

An alternative mechanism by which Dlg could regulate Scrib cortical localization is via physical binding. The conserved colocalization and shared functions of Scrib module proteins has led to propositions that they function as a macromolecular complex. The Dlg GUK domain is the central mediator: it is reported to interact directly with Lgl, and indirectly with Scrib PDZ domains through the protein Gukholder (Gukh); it also binds to the Dlg SH3 domain in an autoinhibitory manner (34–37). We tested the requirement for these interactions *in vivo* by analyzing a *dlg* hypomorphic allele (*dlg*^*v59*^) that removes the GUK domain (21)(**Fig. 2C**). Apicobasal polarity and aPKC localization remained unchanged in central follicle cells mutant for the GUK-deficient allele, as did cortical localization of Lgl (**Fig. 2G,I, Fig. S3D,E**). A partial loss of cortical Scrib localization was seen, although this may be due to reduced levels of mutant Dlg (**Fig. 2H, Fig. S3C**)(21). As with the GUK-deficient *dlg* allele, no polarity defects were seen in cells homozygous for a *scrib* allele that lacks PDZ domains (*scrib*^*4*^) (**Fig. 2J,K**)(38). By contrast, a missense mutation in the Dlg SH3 domain (*dlg*^*m30*^), which does not alter Dlg protein levels or localization, was sufficient to cause mislocalization of Scrib, as well as both Lgl and aPKC, in a manner indistinguishable from null alleles (**Fig. 2L,M, Fig. S3A**)(21). These results reveal a role for the SH3 domain in regulating Scrib localization as well as apical domain antagonism, but show that the GUK domain is not required for epithelial polarity.

A third mechanism that can localize cytosolic proteins to the cortex is via post-translational attachment of lipophilic groups. We performed Acyl-Biotin Exchange (ABE) on larval lysates and found that Scrib::GFP is acylated in *Drosophila* (**Fig. 3A**), consistent with a previous report (39, 40). Recent work has shown that mammalian Scrib is S-palmitoylated on two conserved N-terminal cysteine residues, and this modification is required for Scrib cortical localization and function (41). We generated a Scrib::GFP transgene in which these conserved palmitoylated cysteines are changed to alanine (Scrib^C4AC11A^::GFP). Surprisingly, this protein localizes appropriately to the plasma membrane and rescues *scrib* mutant polarity phenotypes (**Fig. 3B-D**). ABE showed that these mutations are not sufficient to abolish all acylation, suggesting that Scrib can be palmitoylated on additional non-conserved residues (**Fig. 3A**). We then inhibited palmitoyltransferases using chemical and genetic means, and found that both of these approaches failed to impact Scrib localization (**Fig. S4A-C**). Finally, we asked whether Dlg might regulate Scrib through influencing its palmitoylation. However, in *dlg* tissue no change in the acylation of Scrib::GFP could be detected (**Fig. 3A**). Thus, Dlg regulates Scrib membrane localization by a mechanism independent of palmitoylation.

**Figure 3.**
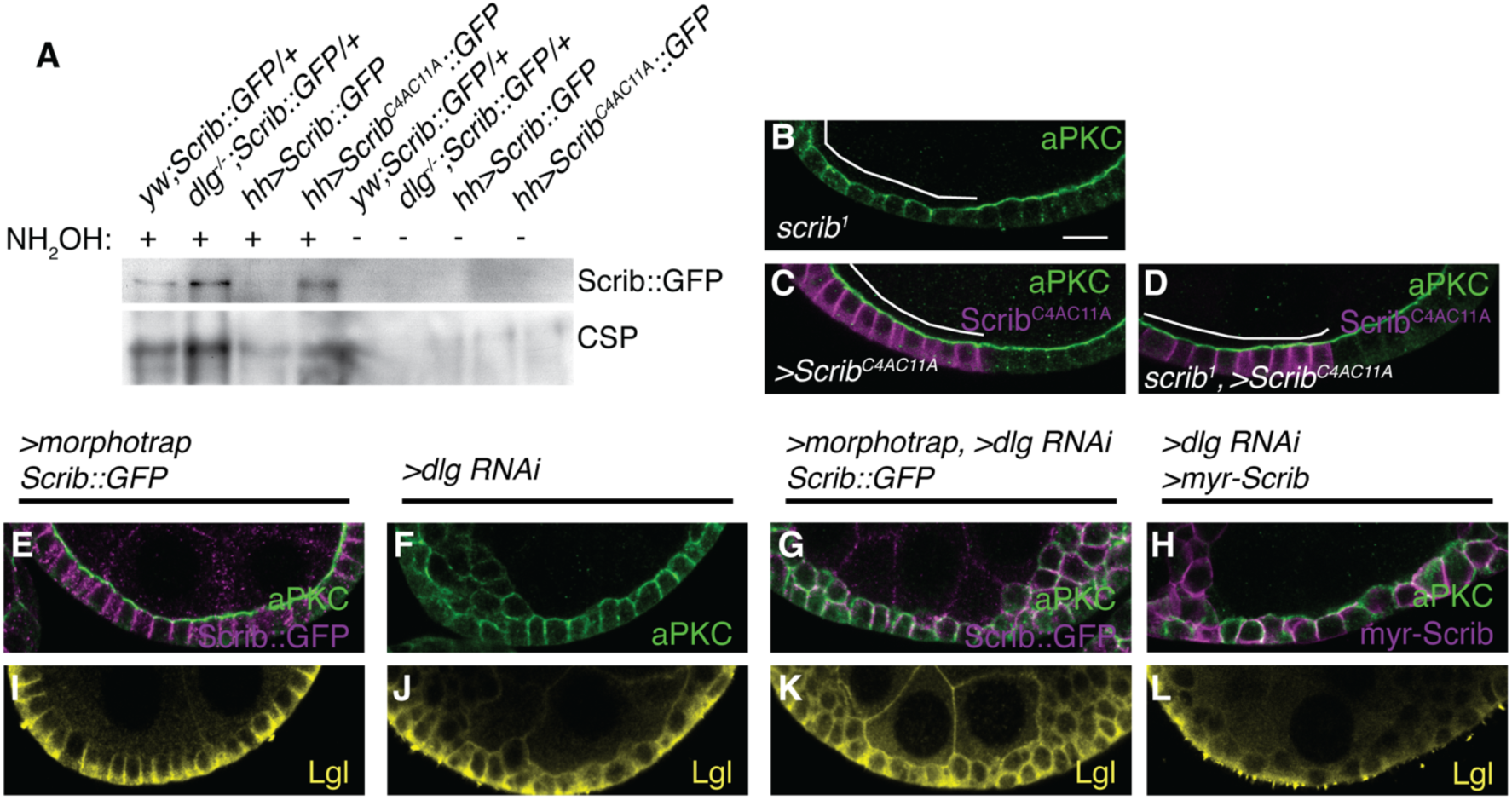
Dlg has Scrib-independent polarity functions. (A) ABE demonstrates that Scrib is palmitoylated *in vivo* in larval tissues, but this is not detectably altered in *dlg* mutant animals. Mutating two conserved cysteines (Scrib^C4AC11A^) does not abolish Scrib palmitoylation. Cysteine String Protein (CSP) serves as a control palmitoylated protein. Non-NH_2_OH treated lanes control for biotinylation specificity to palmitoylated residues. (B-D) Scrib^C4AC11A^ can fully rescue polarity loss in *scrib* mutant follicle cells, and localizes appropriately to the basolateral membrane. Compared to WT and *dlg* RNAi alone (E,F,I,J), membrane tethering Scrib by Morphotrap (G,K) or N-terminal myristoylation signal (H,L) in *dlg* RNAi cells does not rescue aPKC spread or Lgl mislocalization. Scale bars, 10µm.

To test whether cortical Scrib stabilization is the sole function of Dlg in epithelial polarity, we made use of a nanobody-based system for relocalizing GFP-tagged proteins within the cell (42). We tethered Scrib::GFP to the cortex via interactions with a uniformly distributed transmembrane anchor and examined apicobasal polarity in the absence of *dlg.* However, aPKC mislocalized to the lateral membrane and Lgl was displaced to the cytoplasm in these cells, as in cells depleted of *dlg* alone (**Fig. 3E-G,I-K**). As a complementary approach, we generated a constitutively membrane-tethered version of Scrib via attachment of an N-terminal myristoylation sequence. This myr-Scrib transgene was capable of rescuing polarity defects in *scrib* mutant follicle cells (**Fig. S5A-E**). However, in *dlg*-depleted cells expressing myr-Scrib, in which myr-Scrib remains cortical, neither aPKC nor Lgl mislocalization was rescued (**Fig. 3H,L**). From these experiments, we conclude that Dlg has polarity functions independent of Scrib stabilization and that both module components act in parallel to regulate apicobasal protein localization.

### Scrib and Dlg are not regulated by, and do not directly regulate, aPKC

We then examined the relationship between the Scrib module and aPKC. A central feature of this relationship is the exclusion of Lgl from aPKC-containing membrane domains, due to direct phosphorylation; when follicle cells are depleted of *apkc*, Lgl can reach the apical membrane (**Fig. 4A**)(43–45). We asked if Scrib and Dlg also exhibit aPKC-dependent apical exclusion, but found that Dlg and Scrib remain polarized to the basolateral membrane in *apkc*-depleted follicle cells (**Fig. 4B,C**). Dlg and Scrib also remain basolaterally restricted when aPKC is depleted within *lgl* mutant cells (**Fig. 4D,E,G,H**). Additionally, overexpression of a constitutively active form of aPKC (aPKC^ΔN^) does not displace Scrib or Dlg from the plasma membrane (**Fig. 4M**). Thus, polarization of Scrib and Dlg depends on cues independent of aPKC activity.

**Figure 4.**
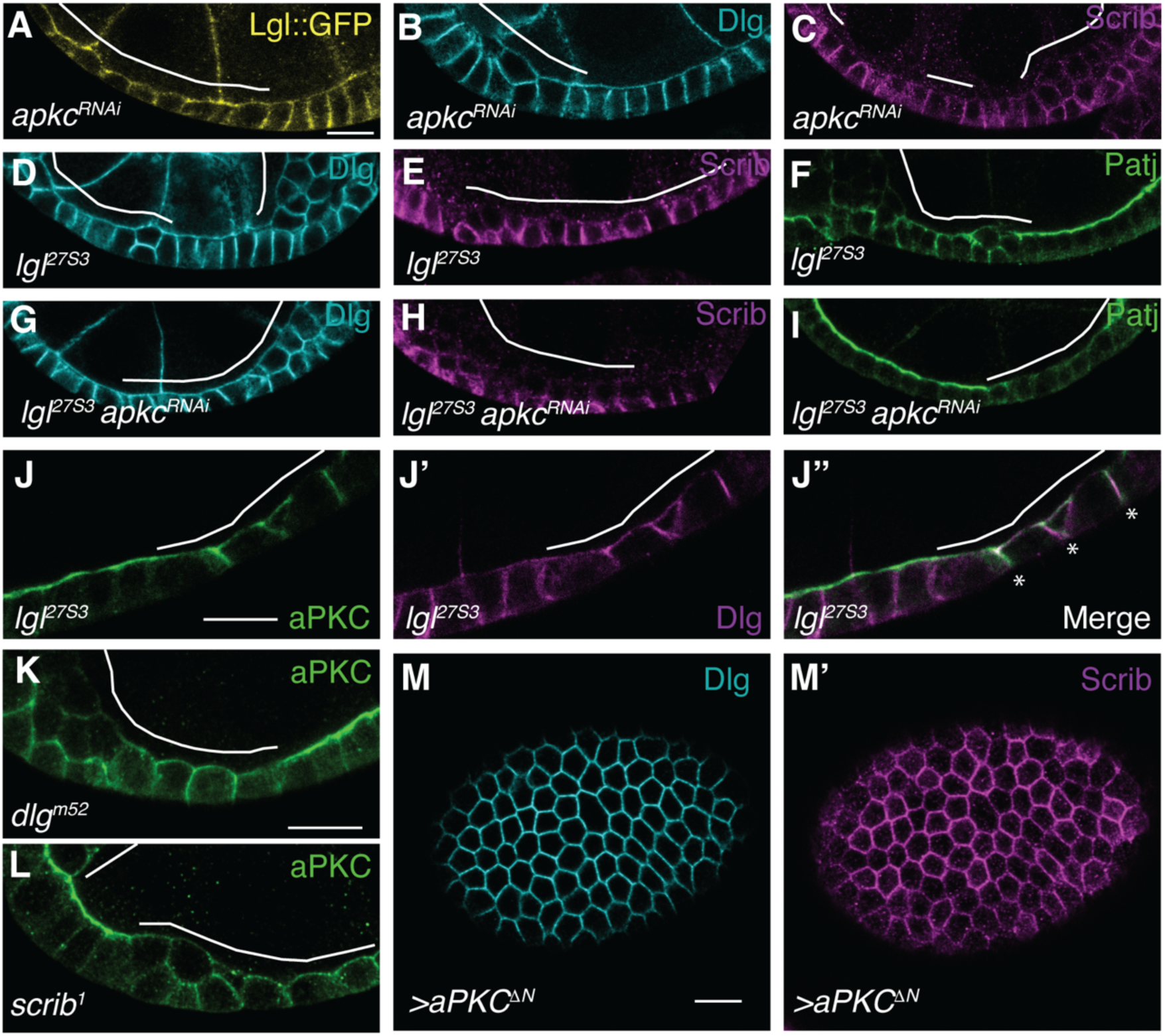
Scrib and Dlg do not directly antagonize aPKC. (A) In *apkc*-depleted follicle cells, Lgl reaches the apical membrane, but Dlg (B) and Scrib (C) remain basolaterally localized. (D,E,G,H) Scrib and Dlg localization is unaffected when apicobasal antagonism is eliminated by codepletion of *apkc* and *lgl.* Laterally mislocalized aPKC in *lgl* mutant cells (F) is active, as it recruits Patj to these sites (I). (J) aPKC spreads along the basolateral membrane in *lgl* mutant cells, where it colocalizes with Dlg (J’,J”) and Scrib (not shown). (K,L) aPKC mislocalization is also seen in *dlg* (K) and *scrib* (L) mutant cells. (M) Expression of a constitutively active aPKC (aPKC^ΔN^) does not displace Scrib or Dlg. Scale bars, 10µm.

The inhibitory relationship between aPKC and Lgl is well established, but it is not known whether Scrib and Dlg might also be direct inhibitors of aPKC (12, 15, 16). Notably, when aPKC mislocalizes laterally in *lgl* mutant cells, it colocalizes with Scrib and Dlg, which are not displaced (**Fig. 4J**). This lateral aPKC is active because it can recruit Patj, an aPKC-dependent apical protein to ectopic sites (**Fig. 4F,I**)(9). These results suggest that Lgl alone has intrinsic ability to inhibit aPKC, and that the aPKC mislocalization seen in *scrib* and *dlg* mutant cells (**Fig. 4K,L**) reflects a weakening of Lgl inhibitory activity in the absence of either Scrib or Dlg.

### Scrib and Dlg are both required to stabilize and enable Lgl activity

If Scrib and Dlg do not directly inhibit aPKC, how do they participate in apicobasal antagonism? FRAP measurements of Lgl::GFP show that in *dlg* and *scrib-*depleted follicle cells, a clear increase in recovery kinetics and decrease of the mobile fraction compared to WT is seen (**Fig. 5A,B**). Whereas Lgl::GFP becomes cytoplasmic in *dlg* RNAi cells, an endogenously expressed, non-phosphorylatable Lgl fusion protein (Lgl^S5A^::GFP) remains cortically associated in *dlg* RNAi cells (**Fig. 5C,D**)(13). Moreover, co-depleting aPKC in *dlg* RNAi cells restores Lgl cortical localization (**Fig. 5E,F**). These results are consistent with dynamic exchange of Lgl between an hypophosphorylated membrane-associated pool and an aPKC-hyperphosphorylated cytoplasmic pool, and suggest that Scrib and Dlg stabilize the former.

**Figure 5.**
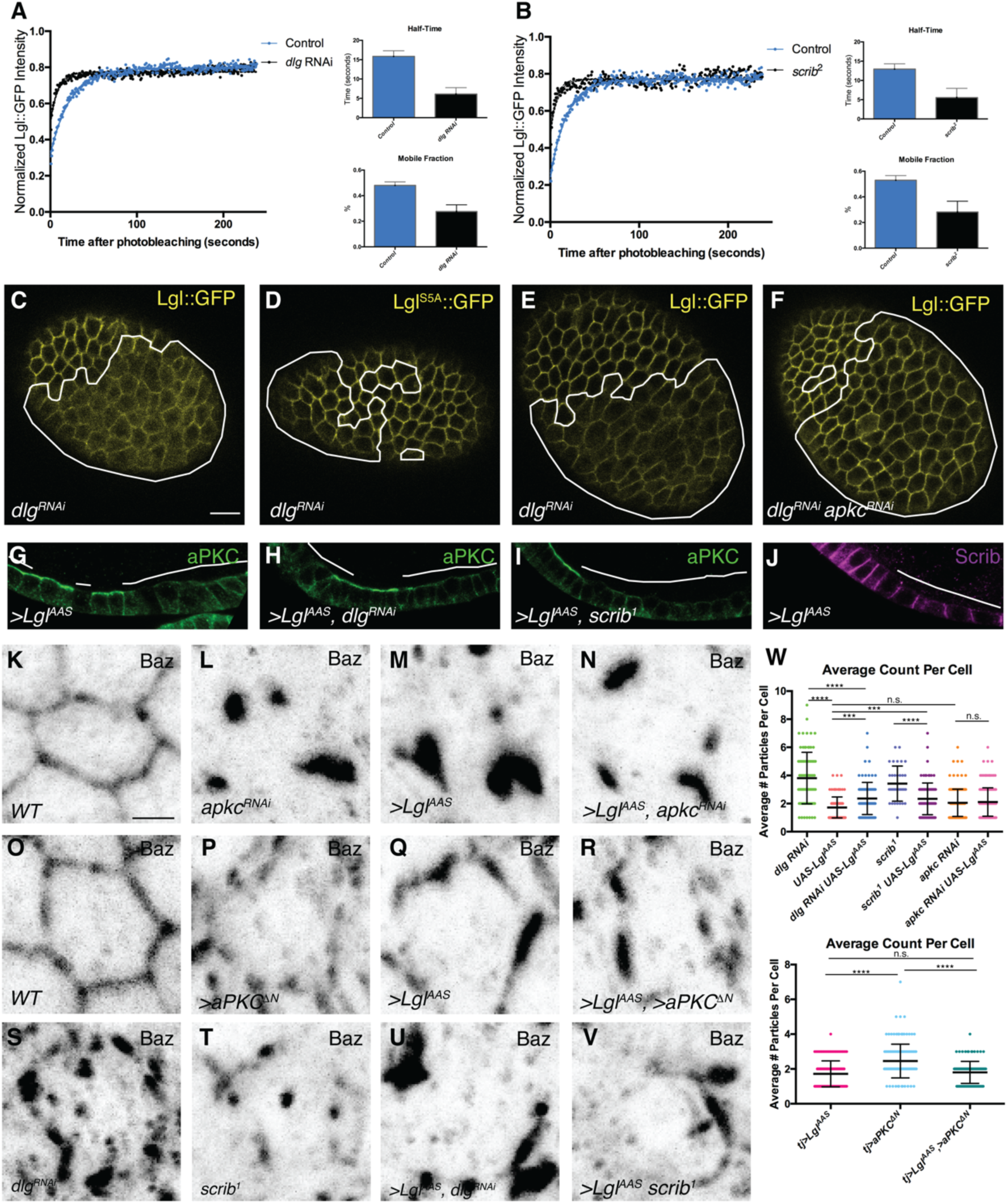
Scrib and Dlg support basolateral Lgl activity. (A,B) In both *scrib* and *dlg* mutant follicle cells, Lgl::GFP shows increased FRAP kinetics compared to WT. (C,D) In *dlg* RNAi cells, Lgl::GFP is displaced to the cytoplasm but non-phosphorylatable Lgl^S5A^::GFP remains cortical. (E,F) Co-depletion of *apkc* rescues Lgl localization in *dlg-*depleted cell. (G) Lgl^AAS^ expression causes loss of apical aPKC. (H,I) Apical aPKC depletion by Lgl^AAS^ persists in the absence of Dlg and Scrib. (J) Lgl^AAS^ is not sufficient to create an ectopic basolateral membrane, as it fails to recruit Scrib apically. (K-M) Baz forms several aggregates in each Lgl^AAS^-expressing cell, similar to *apkc* depleted cells. (M,N) Depletion of *apkc* in Lgl^AAS^-expressing cells does not modify the Baz phenotype. Compared to WT (O), expressing of a constitutively active aPKC (P) causes a Baz phenotype similar to *dlg* (S) or *scrib* loss-of-function (T). (Q,R) Co-expression of Lgl^AAS^ causes a phenotype that resembles Lgl^AAS^ alone. In *dlg*-depleted (S) or *scrib* mutant (T) cells, Baz localizes to more frequent, fragmented puncta. Expression of Lgl^AAS^ in *dlg*-depleted cells (U) or *scrib* mutant cells (V) reduces Baz particle number. (W) Quantification of Baz phenotypes in K-V. One-way ANOVA with Tukey’s multiple comparisons test. K-V show single cell maximum intensity projections. Scale bars, 10µm in C-J, 2µm in K-V. Error bars in A-B represent 95% confidence intervals, error bars in W represent S.D.

Cortical association of Lgl depends on interactions between polar phospholipids and charged residues within the Lgl PBM (13, 14). PIP2 and PIP3 show apicobasally polarized distributions in epithelial cells of *Drosophila* as well as vertebrates, raising the possibility that Dlg and Scrib could regulate Lgl function by altering the distribution of PIP species at the basolateral membrane (46, 47). However, using reporters for PIP2 and PIP3, we did not detect differences in their distribution or levels in *dlg* or *scrib* mutant cells (**Fig. S6**)(48, 49).

An alternative mechanism by which Scrib and Dlg could ensure antagonism of apical identity is by simply promoting Lgl cortical localization. We therefore tested whether membrane localization of Lgl was sufficient to bypass loss of *scrib* or *dlg* function in follicle epithelia. However, in our hands overexpression of a constitutively membrane-tethered Lgl (myr-Lgl) did not alter polarity defects in *scrib-* or *dlg*-depleted follicle cells, nor did it cause polarity defects in WT follicle cells (**Fig. S5F-J**)(44). By contrast, a mutant Lgl protein with only the most C-terminal aPKC phosphorylation site present (Lgl^S656A,S660A^, hereafter Lgl^AAS^) was suggested to be a dominant inhibitor of aPKC (43). We confirmed that Lgl^AAS^ expression in otherwise WT follicle cells causes dominant phenotypes, including multilayering and loss of apical aPKC staining (**Fig. 5G**). We note that although Lgl^AAS^ localizes uniformly to the cortex including the apical domain and can displace aPKC, it cannot establish an ectopic basolateral domain at the former apical site, as it does not recruit Scrib (**Fig. 5J**).

To determine whether Lgl^AAS^ is a bona fide aPKC inhibitor, we compared the phenotype of Lgl^AAS^ -expressing cells with *apkc* RNAi-expressing cells, using Bazooka (Baz, *Drosophila* Par-3) localization as a phenotypic readout (7, 11). Baz is an aPKC substrate, and preventing phosphorylation via *apkc* depletion or expression of non-phosphorylatable Baz results in formation of several large aggregates in the cell (**Fig. 5L**)(8–11, 50–52). Interestingly, Lgl^AAS^ also induced Baz aggregates (**Fig. 5M**), and co-depletion of *apkc* did not modify the phenotype (**Fig. 5N**). Furthermore, while expression of an activated form of aPKC caused Baz to localize in a larger number of fragmented puncta, similar to those described previously in basolateral mutants (**Fig. 5O,P vs. S,T**)(20, 53), coexpression of Lgl^AAS^ resulted in aggregates indistinguishable from those caused by expression of Lgl^AAS^ alone (**Fig. 5Q,R,W**). These data are consistent with a model in which Lgl^AAS^ dominantly affects apicobasal polarity by inhibiting aPKC.

We then asked whether the dominant effects of Lgl^AAS^ depend on Dlg or Scrib activity. In *dlg* RNAi or *scrib* mutant cells, Lgl^AAS^ retained the ability to create several Baz aggregates, although an increased number suggested incomplete epistasis (**Fig. 5S-W**). Coexpression of Lgl^AAS^ also reduced the lateral expansion of aPKC seen in cells depleted of either *dlg* or *scrib* (**Fig. 4K, L vs 5G-I**). These results suggest that many apical-inhibiting effects of Lgl^AAS^ do not strictly depend on Scrib or Dlg.

### The entire Scrib module is necessary but not sufficient for basolateral polarity

The fact that the activity of WT Lgl but not Lgl^AAS^ requires Scrib and Dlg suggests that Scrib and Dlg could enhance Lgl’s ability to antagonize aPKC at the basolateral membrane, perhaps by protecting Lgl from aPKC phosphorylation. A model where both Scrib and Dlg are required would be consistent with the inability of either single protein to bypass loss of the other (**Fig. 1R-W**). To test if ectopic apical localization of Scrib and Dlg together would therefore allow Lgl to inhibit aPKC, we used a combination of apical domain-specific nanbody tethering and overexpression to simultaneously mislocalize one, two, or all three Scrib module proteins (42). However, despite robust colocalization at the apical cell surface, no effects were seen in any case on aPKC, apicobasal polarity, or epithelial architecture (**Fig. 6A-J**). We conclude that, despite the necessity for each component in basolateral domain identity, even the entire Scrib module together is not sufficient to inhibit apical polarity determinants.

**Figure 6.**
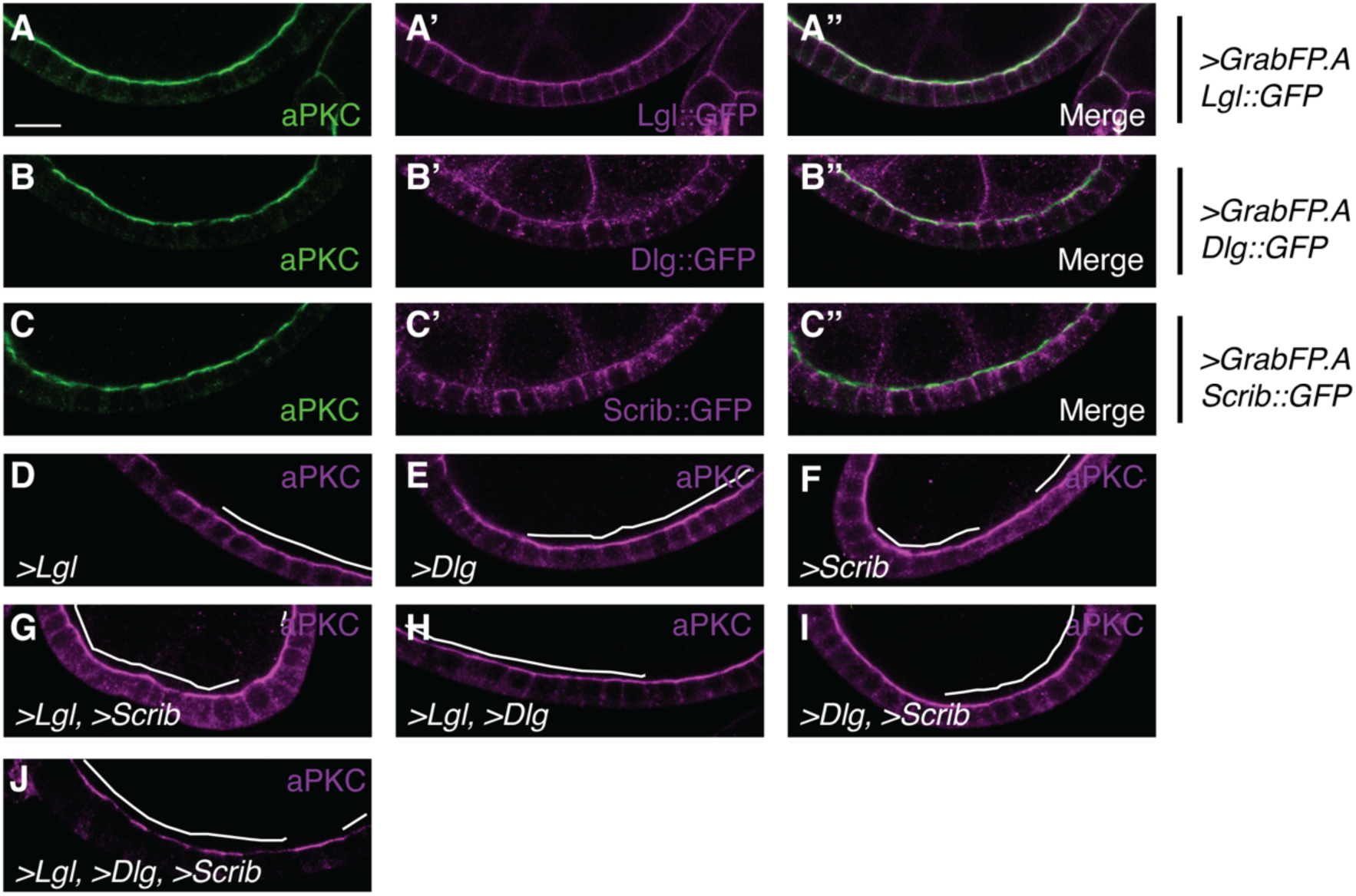
The entire Scrib module is not sufficient to establish basolateral polarity. Ectopic apical localization of Lgl (A), Dlg (B) or Scrib (C) using apical GrabFP has no effect on epithelial polarity. WT Scrib, Dlg and Lgl are not sufficient to disrupt the apical domain when overexpressed singly (D-F), in pairs (G-I), or as a holo-module (J). Scale bars, 10µm.

## DISCUSSION

Despite being central regulators of cell polarity in numerous systems, the mechanisms of Scrib module activity have remained obscure. Our work highlights previously unappreciated specificity in these activities, and begins to define the molecular functions of Scrib, Dlg and Lgl. We did not observe phenotypic enhancement in double mutant follicle cells compared to single mutants, which together with the complete penetrance of single mutant phenotypes suggests full codependence of function rather than functional overlap. Moreover, we were unable to bypass Scrib module mutants in any combination by overexpression, consistent with unique roles for each protein. Thus, while Scrib, Dlg and Lgl act in a common “basolateral polarity” pathway, they each contribute distinct functions to give rise to the basolateral membrane.

Cell polarity is particularly evident at the plasma membrane, and most polarity regulators act at the cell cortex. A key question in the field therefore has been the mechanisms that allow cortical localization of the Scrib module and Par complex proteins, which exhibit no classical membrane association domains (50). We find a simple linear hierarchy for cortical localization in the follicle that places Dlg most upstream, and contrasts with that recently described in the adult midgut, where Scrib appears to be most upstream (26, 28). Our work highlights the requirement of Dlg for Scrib localization, and provides insight into the mechanism, in part by ruling out previous models. One model involves a direct physical interaction, mediated by the Scrib PDZ domains and Dlg GUK domain (23, 24, 35). However, our *in vivo* analyses show that follicle cells mutant for alleles lacking either of these domains have normal polarity; these results are supported by data from imaginal discs (21, 38, 51). In contrast, we show that the SH3 domain is critical for Scrib cortical localization as well as polarity (51). The Dlg SH3 and GUK domains engage in an intramolecular ‘autoinhibitory’ interaction that negatively regulates binding of partners such as Gukh, CASK and CRIPT (37, 52–58). The dispensability of the GUK domain provides evidence against an essential role for this mode of regulation in epithelial polarity, and highlight the value of investigating the GUK-independent function of the Dlg SH3.

We also exclude a second mechanism of Scrib cortical association. Mammalian Scrib is S-palmitoylated, and this modification is required for both cortical localization and function (41). As *Drosophila* Scrib was also recently shown to be palmitoylated, an appealing model would involve Dlg regulating this post-translational modification (40). However, we could detect no changes to Scrib palmitoylation in a *dlg* mutant, and chemically or genetically inhibiting *Drosophila* palmitoyltransferases also had no effect on Scrib localization. Surprisingly, palmitoylated Scrib is incapable of reaching the cortex in *dlg* mutants. While a constitutively myristoylated Scrib can bypass this requirement for localization, it is nevertheless insufficient to support polarity in the absence of Dlg. These results indicate that Dlg has Scrib-independent functions and that Dlg may regulate additional basolateral activities, perhaps by scaffolding as yet unidentified factors.

Lgl’s role as an aPKC inhibitor is well-characterized, but how Scrib and Dlg influence this antagonism is not understood. Our data show that Scrib and Dlg maintain cortical Lgl by regulating its phosphorylation by aPKC, but also suggest that this is not via direct inhibition of aPKC kinase activity or intrinsic antagonism of aPKC localization. Instead, they are consistent with models in which Scrib and Dlg regulate the three specific aPKC-targeted residues in Lgl. Previous work has demonstrated that these phosphorylated serines (656, 660, 664) are neither functionally nor kinetically equivalent, and a recent model proposes that S664 is required for basolateral polarization by mediating a phosphorylation-dependent interaction with the Dlg GUK domain (34, 43, 59, 60). Beyond the dispensability of the GUK domain, the ability of Lgl^AAS^ to inhibit aPKC largely independently of Scrib and Dlg argues against this model. Moreover, only Lgl^AAS^ among the phospho-mutants can dominantly inhibit aPKC, while WT Lgl can do the same only if Scrib and Dlg are present. Together, these results suggest that S656 is the critical inhibitory residue whose phosphorylation must be limited to enable Lgl’s activity. We favor a model in which Scrib and Dlg ‘protect’ Lgl by limiting phosphorylation of this site, thus tipping the inhibitory balance to allow Lgl to inhibit aPKC instead and establish the basolateral membrane.

What mechanism could underlie Scrib and Dlg protection of Lgl? One mechanism could involve generating a high phospholipid charge density at the basolateral membrane, which has been shown to desensitize Lgl to aPKC phosphorylation *in vitro* (61). However, our data do not find evidence for regulation of phosphoinositides by Scrib and Dlg. A second possibility is that Scrib and Dlg could scaffold an additional factor, such as PP1 phosphatase, which counteracts aPKC phosphorylation of Lgl (62). Alternative mechanisms include those suggested by recent work on PAR-1 and PAR-2 in *C. elegans* zygotes, a circuit with several parallels to the Scrib module (63–65). In this system, PAR-2 protects PAR-1 at the cortex by shielding it from aPKC phosphorylation through physical interaction-dependent and -independent mechanisms (63). By analogy, binding with Scrib and/or Dlg could allosterically regulate Lgl to prevent phosphorylation, although we have ruled out Lgl-Scrib and Lgl-Dlg interactions documented in the literature (34, 66). Scrib or Dlg might also act as a “decoy substrate” for aPKC, as PAR-2 does in PAR-1 protection (63). Indeed, Scrib is phosphorylated on at least 13 residues in *Drosophila* embryos, though the functional relevance of this is not yet known (67).

Overall, our work highlights the multifaceted nature of Scrib module function. The failure to bypass Scrib module mutants by transgenic supply of any single or double combination of other module components, including several that were constitutively membrane-tethered, suggests that every member contributes a specific activity to polarity. Nevertheless, even the simultaneous ectopic localization of all three Scrib module proteins was insufficient to disrupt the apical domain. This insufficiency in basolateral specification may reflect an inability of apical Scrib and Dlg to protect Lgl from aPKC phosphorylation, perhaps due to the distinct molecular composition of the apical and basolateral domains. This supports the idea that in addition to intrinsic activity via Lgl, the Scrib module must recruit or activate additional, as yet unidentified effectors in basolateral polarity establishment. The independent as well as cooperative activities of the Scrib module delineated here demonstrate previously unappreciated complexity in the determination of basolateral polarity and set the stage for future mechanistic studies of Scrib module function.

## MATERIALS AND METHODS

Mutant and overexpression analyses employed *hsFLP, GR1-GAL4 UAS-FLP* or *traffic jam-GAL4. UAS-myr-Scrib::V5* was generated by appending the N-terminal myristoylation signal from Src42A and C-terminal V5 tag to the Scrib A2 cDNA, and *UASp-Scrib*^*C4AC11A*^::*GFP* was generated via site-directed mutagenesis. Acyl-Biotin Exchange was performed by modifying published protocols (39, 68), using anterior L3 larval lysates; biotinylated protein was purified using magnetic beads and analyzed by western blot. Images were acquired using Zeiss LSM700 or LSM780 laser scanning confocal microscopes with LD C-Apochromat 40x/NA1.1 W or Plan Apochromat 63x/NA1.4 oil objectives. Image processing and quantification was performed using Fiji software (69); for significance in statistical tests: n.s.=p>0.05, *=p<0.05, **=p<0.01, ***=p<0.001 and ****=p<0.0001. FRAP experiments were performed as previously described, and processed and analyzed using Fiji and Graphpad Prism (69, 70). Baz particles were quantified by the Analyze Particles function and the FeatureJ plugin. Details are provided in the *SI Appendix, Materials and Methods.*

## Acknowledgments

We thank E. Morais de Sá, D. Bergstralh, B. Thompson, V. Budnik, T. Schupbach, R. Davis, Y. Hong, F. Matsuzaki and I. Hariharan for generously providing fly stocks and reagents and C. Brownlee and K. Strassburger for providing technical expertise for the ABE experiments. We thank Laura Mathies for cloning the modified Scrib constructs. Imaging on the LSM780 was performed at the CRL Molecular Imaging Center, supported by NSF DBI-1041078. We are grateful to G. de Vreede, M. Peifer and T. Bonello, K. Prehoda, M. Kitaoka and the Bilder laboratory for helpful discussion. This work was supported by NIH grants GM068675, GM090150 and GM130388 to D.B.

## Figures and Tables

**Supplemental Figure 1.**
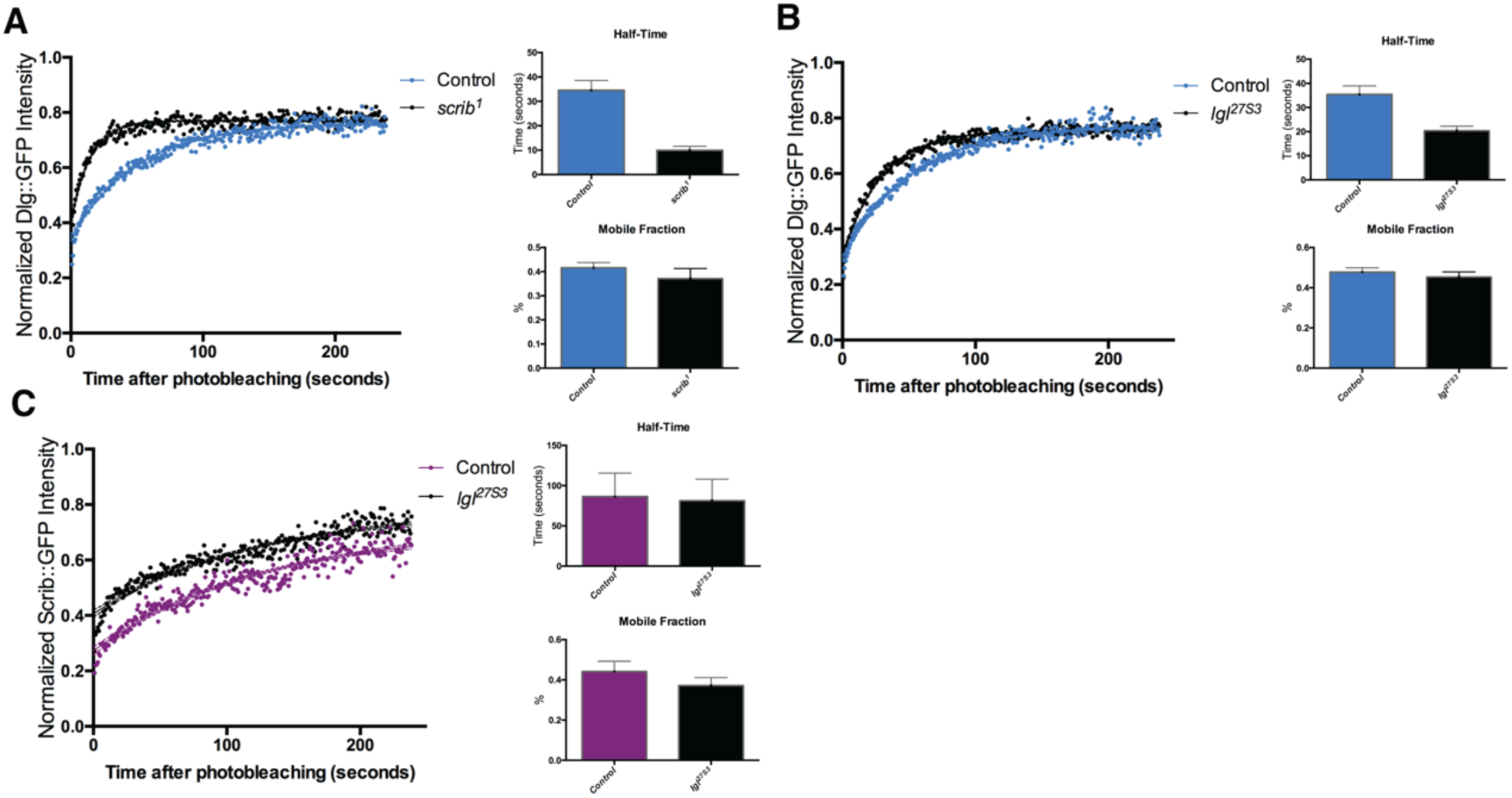
(A) Dlg::GFP shows increased FRAP recovery kinetics in *scrib* mutant follicle cells compared to controls. (B) Dlg::GFP increased FRAP kinetics are slightly increased in *lgl* mutant follicle cells. (C) Scrib::GFP does not show altered FRAP mobility in *lgl* mutant cells. Error bars represent 95% confidence intervals.

**Supplemental Figure 2.**
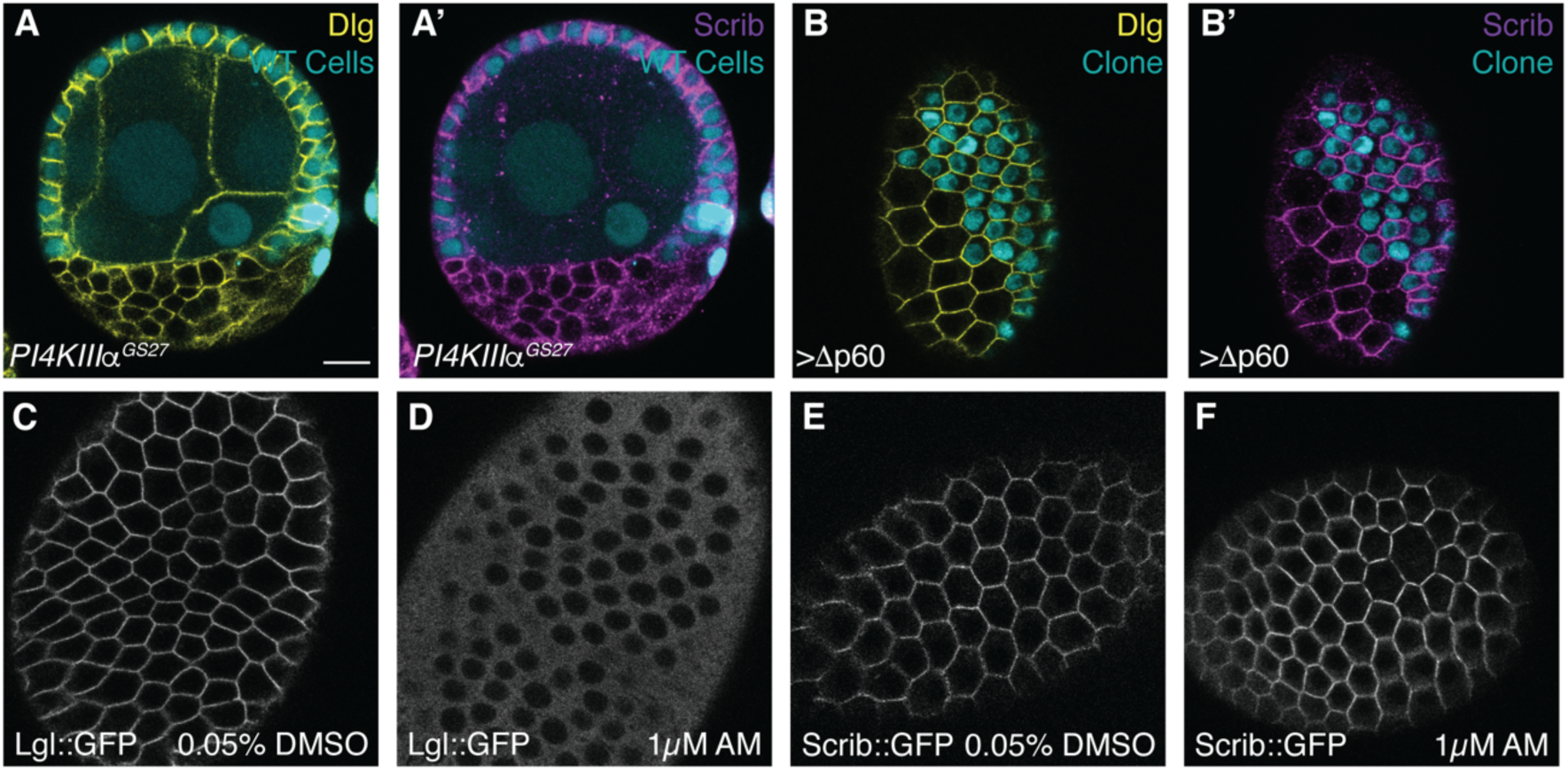
(A) PIP2 depletion by *PI4KIIIα* mutation does not displace Scrib or Dlg membrane localization. (B) PIP3 depletion by expressing dominant negative PI3K (Δp60) does not displace Scrib or Dlg membrane localization. (C,D) ATP depletion by antimycin A (AM) treatment causes Lgl::GFP to become cytoplasmic in follicle cells. (E,F) Scrib::GFP localization is not affected by AM treatment. Scale bars, 10µm.

**Supplemental Figure 3.**
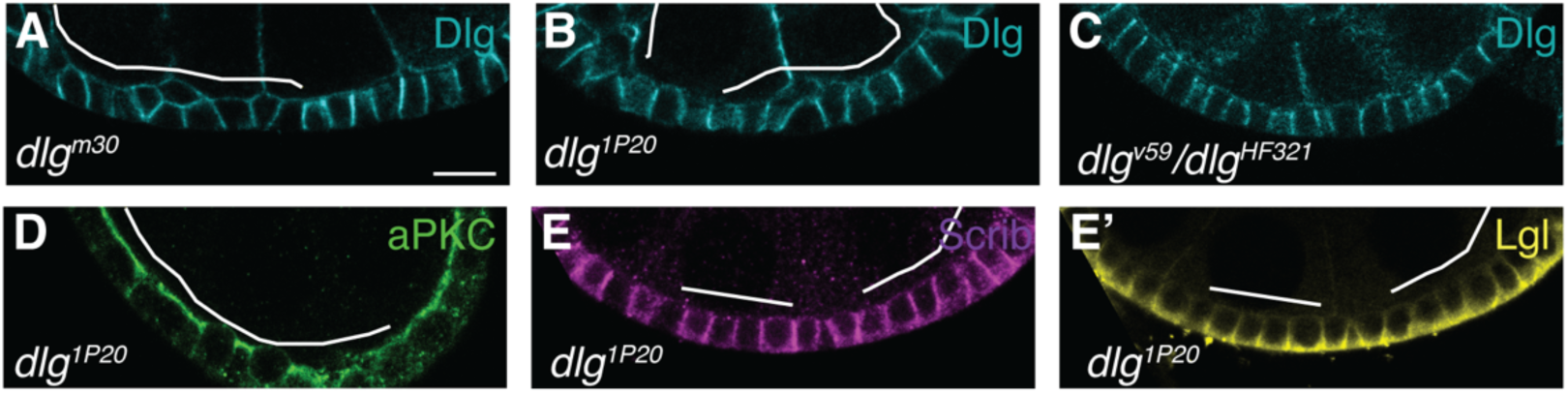
(A-C) Dlg protein is stable and cortically localized in SH3 mutant (A) and GUK-truncated (B,C) mutant follicle cells. (C) Dlg protein levels are decreased in *dlg*^*v59*^ mutant cells, as previously described (21). Follicle cell clones homozygous for a second, less severe, GUK-truncating *dlg* allele shows normal aPKC (D) and Lgl (E’) localization. (E) Scrib localization is also not affected. Scale bars, 10µm.

**Supplemental Figure 4.**
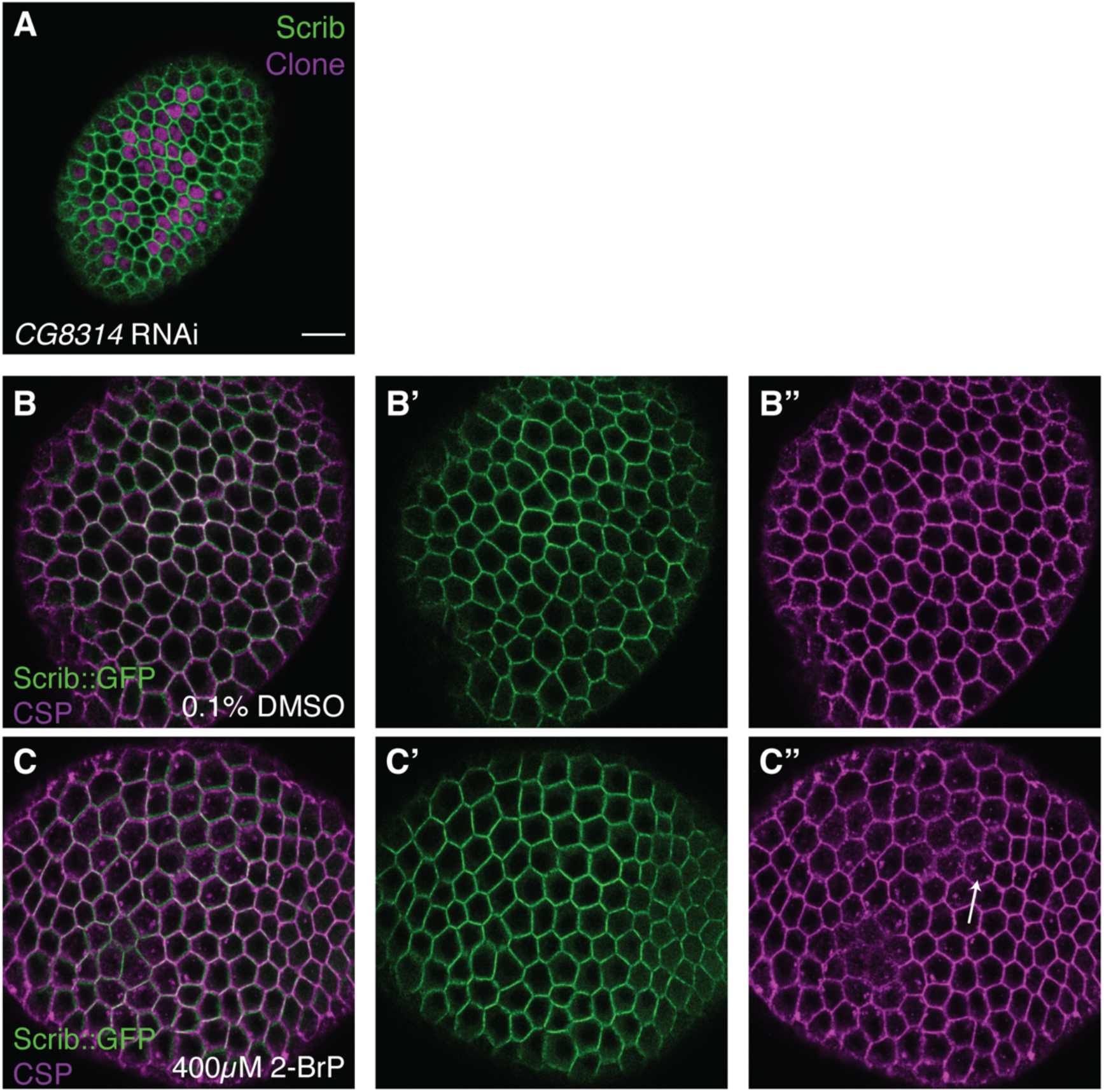
(A) Reducing the levels of CG8314, the homolog to the human Scrib palmitoyltransferase ZDHHC7 does not affect Scrib localization in follicle cells. (B,C) Chemical inhibition of ZDHHC palmitoyltransferases by 2-Bromopalmitate (2-BrP) treatment in *ex vivo* cultured follicles affects a control palmitoylated protein, CSP (B”,C”, arrow), but does not affect Scrib localization (B’,C’). Scale bars, 10µm.

**Supplemental Figure 5.**
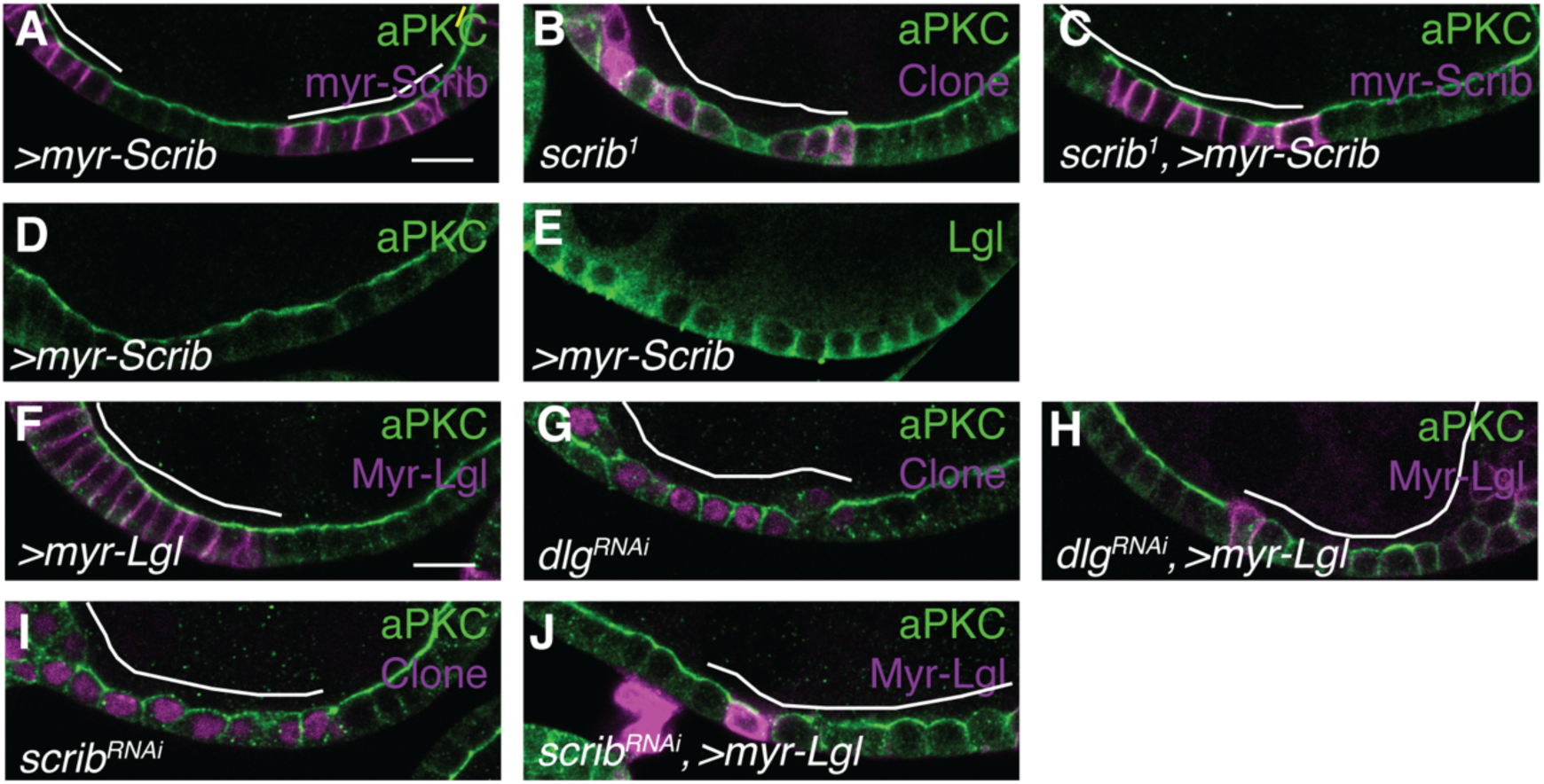
(A) Constitutively membrane-associated myr-Scrib is localized to the membrane in WT cells. (B, C) myr-Scrib rescues polarity loss in *scrib* mutant cells. (D,E) myr-Scrib expression does not disrupt aPKC (D) or Lgl (E) localization in WT cells. (F) Expression of constitutively membrane-associated myr-Lgl does not cause polarity defects in WT follicle cells. (G-J) myr-Lgl also does not rescue the polarity defects of *dlg* RNAi (G,H) or *scrib* RNAi (I,J) cells. Scale bars, 10µm.

**Supplemental Figure 6.**
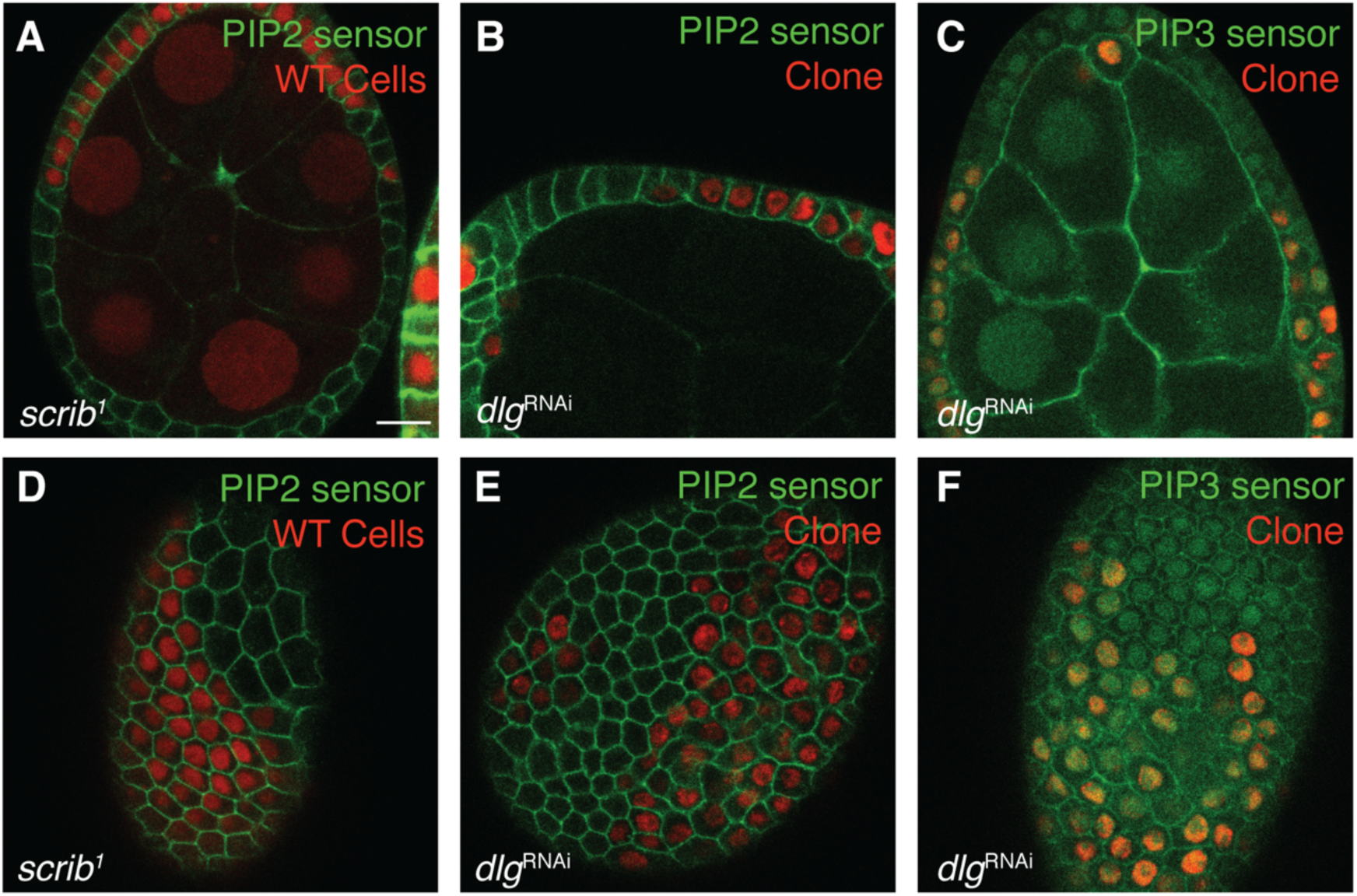
(A,D) *scrib* loss of function does not alter PIP2 levels or localization. (B,E) *dlg* loss of function also does not alter PIP2 levels or localization. (C,F) *dlg* loss of function does not alter PIP3 levels or localization. Scale bars, 10µm.

## Supporting Information

### SI Materials and Methods

#### Fly stocks and genetics

*Drosophila* stocks were raised on cornmeal molasses food at 25°C. *yw* was used as the WT control. Mutant alleles and transgenic lines used are listed in **Table 1**. Mutant follicle cell clones were generated using either *hsFLP* or *GR1-GAL4 UAS-FLP*. Follicle cell MARCM clones were generated using *hsFLP*. For *hsFLP*-induced clones, larvae were heat shocked for 1 hour on two consecutive days starting 120 hours after egg deposition (AED). For clonal GAL4 expression using *hsFLP*, larvae were heat shocked once for 13 minutes 120 hours AED. For all clones, adult females were fed with yeast and dissected 3 days after eclosion. Because *dlg*^*v59*^ is on a chromosomal inversion, it cannot be used with FRT-based recombination, so it was analyzed in trans to *dlg*^*HF321*^, which does not produce protein at 29°C. Pan-follicle cell expression was induced in adults using *traffic jam-GAL4 (tj-GAL4)* and temperature-sensitive GAL80; these were fed yeast for 2 days before shifting to 29°C for 4 days.

**Table 1.**
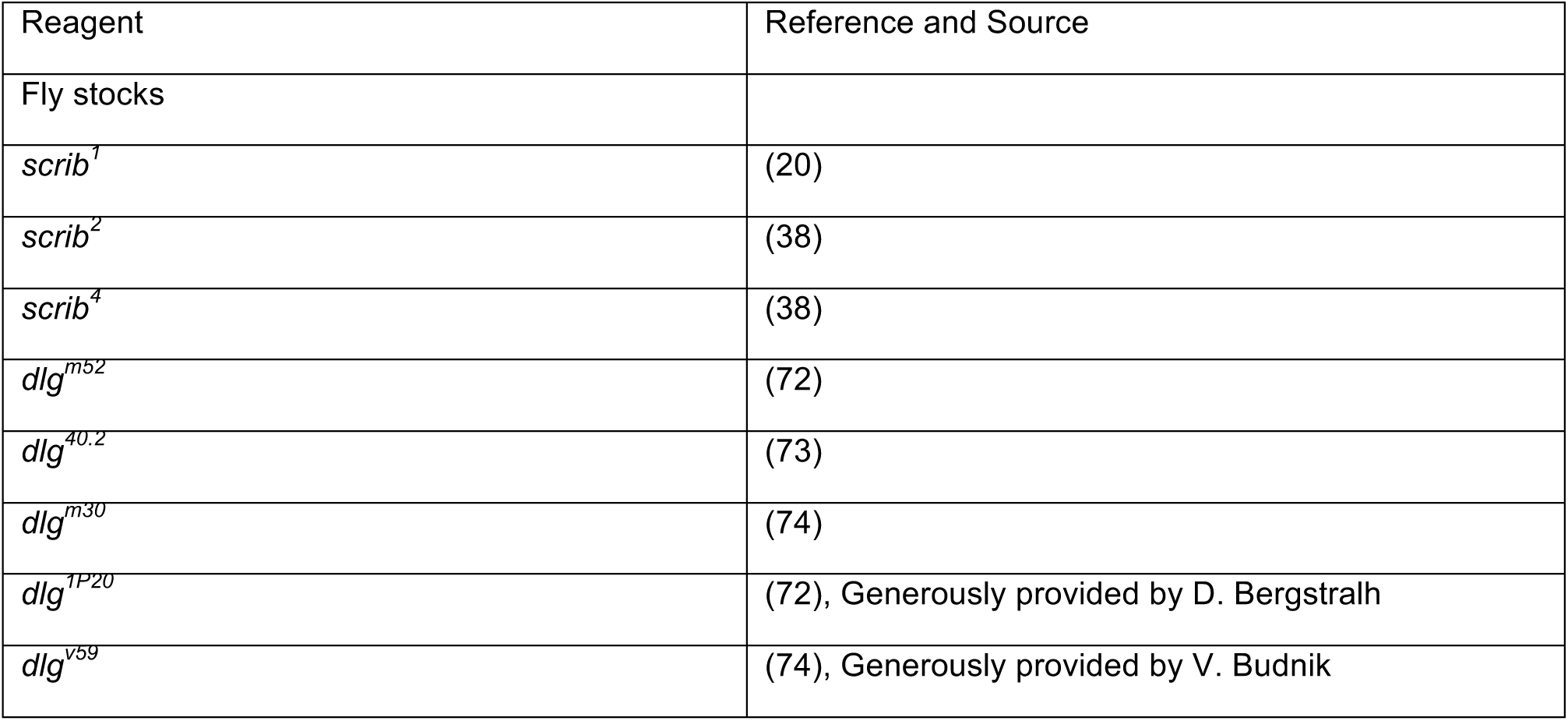
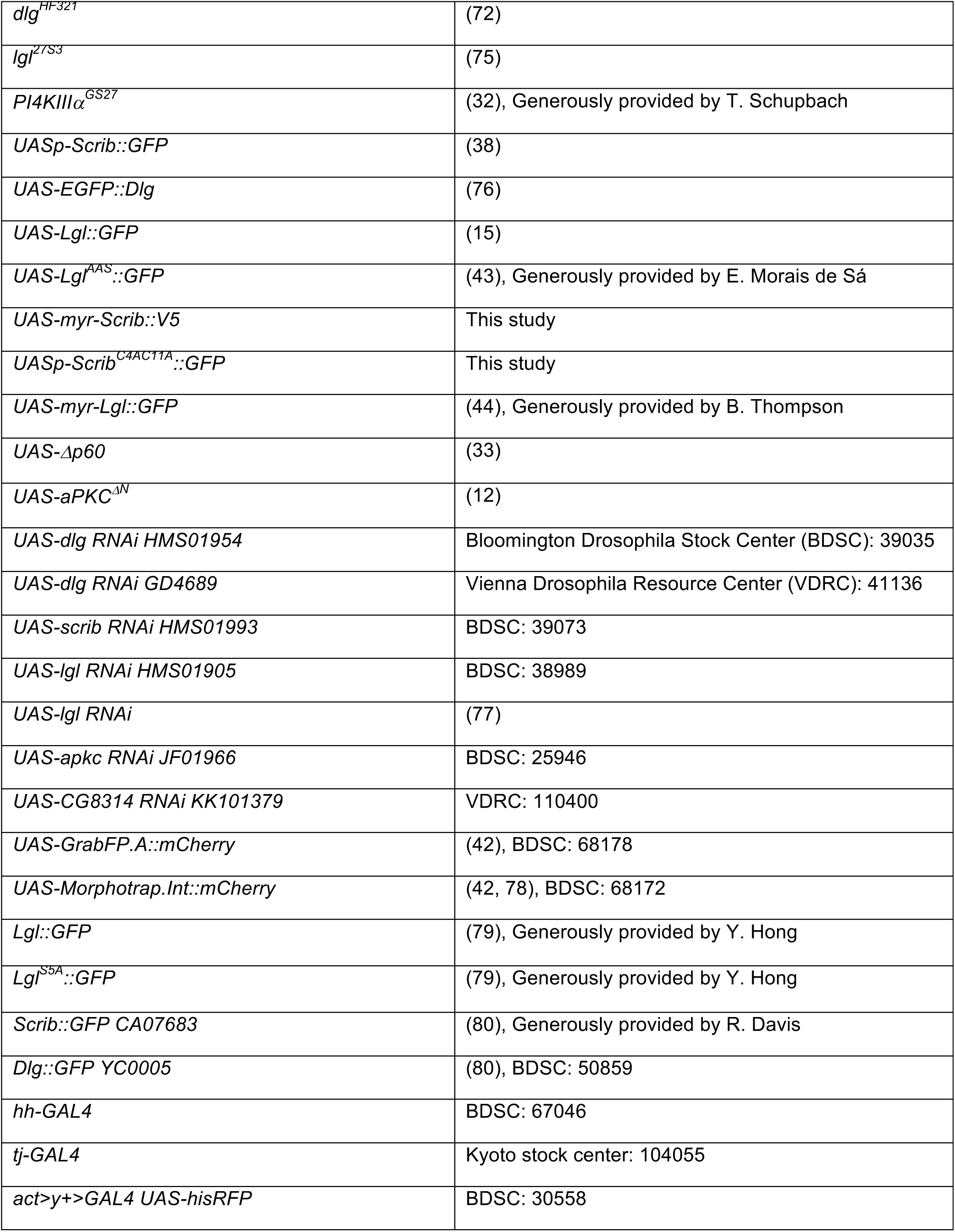
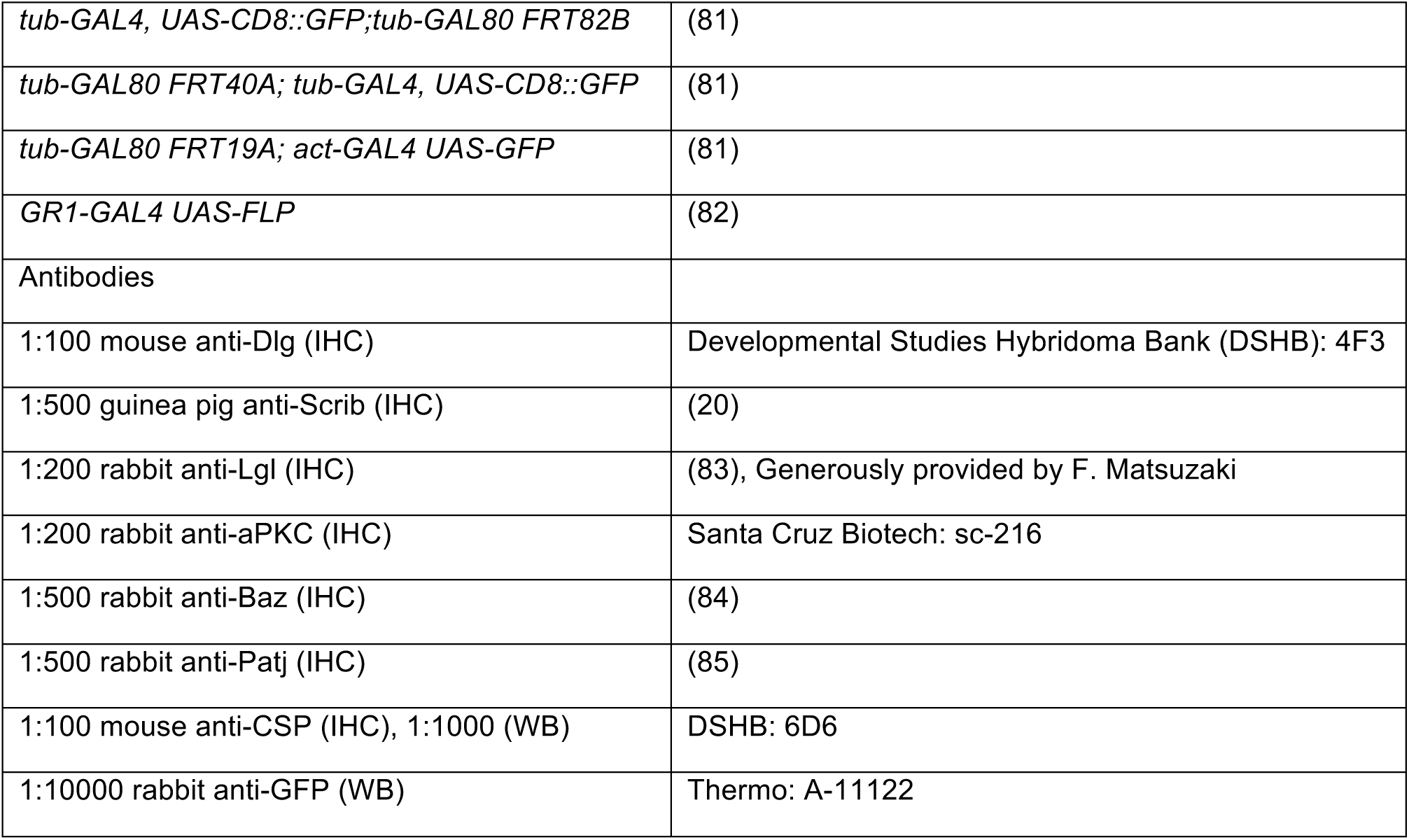
Key resources.

To generate *UAS-myr-Scrib::V5*, the N-terminal myristoylation signal from Src42A (ATGGGTAACTGCCTCACCACACAGAAGGGCGAACCCGACAAGCCCGCA) and C-terminal V5 tag (GGTAAGCCCATTCCAAACCCACTTCTCGGTCTGGATAGCACA) were synthesized as gBlocks with overlap to the pUASTattB backbone and Scrib CDS. The Scrib A2 CDS was amplified from pBS-ScribA2 using primers GTCCTGGGACTCAACGACAT and CGGAGTGGGTTTGGCTCTAA. These fragments were cloned into the EcoRI/XbaI linearized pUASTattB vector using a High Fidelity Gibson Assembly kit (NEB). The *UASp-Scrib*^*C4AC11A*^::*GFP* construct was generated by mutating Scrib C4 and C11 in the pBS-ScribA2 vector using the Q5 site directed mutagenesis kit (NEB) and primers TTCAAGGGCGCCAACCGGCAGGTGGAGTTCG and GATGGGAATGGCCTTGAACATGCTCGTCTTC. Following sequence verification, mutant pBS-ScribA2 was digested with KpnI and EcoRV and this fragment was cloned into the pUASp backbone. *UAS-myr-Scrib::V5* was targeted to the attP40 landing site and *UASp-Scrib*^*C4AC11A*^::*GFP* was inserted by P element-mediated transformation through embryo injections performed by Bestgene, Inc.

#### Immunofluorescence and microscopy

Follicles were dissected in Schneider’s medium containing 15% FBS and fixed with 4% PFA in PBS for 20 minutes. Follicles were stained in PBS containing 0.1% Triton X-100, 1% BSA and 4% NGS overnight at 4°C. Primary antibodies and dilutions are listed in **Table 1**. Following secondary antibody incubation for 2 hours at room temperature, tissue was mounted in glycerol-based antifade (Invitrogen).

Images were acquired using an inverted Zeiss LSM700 or upright LSM780 laser scanning confocal microscopes with LD C-Apochromat 40x/NA 1.1 W or Plan apochromat 63x/NA 1.4 oil objectives. For each experiment, tissue from at least 5 females was analyzed and at least 10 ovarioles and 20 individual follicles were examined.

#### Fluorescence Recovery After Photobleaching (FRAP) experiments

FRAP experiments were performed as previously described (68). Briefly, follicles were dissected in media as above, supplemented with 2% human insulin (Sigma) and embedded in 0.5% low melting agarose in a glass bottom dish. Imaging was performed on an inverted Zeiss LSM700 using a LD C-Apochromat 40x/NA 1.1 W objective. Images were acquired continuously with resolution of 512 x 269 pixels and scan time of 821.67 msec. 10 pre-bleach images were acquired before an elliptical ROI covering one en face cell boundary was bleached twice with a 488nm 10mW laser at 70% power with pixel dwell of 100.85 µsec. Intensities of the bleached region, reference region and background were manually measured using Fiji(69). Background and imaging-dependent photobleaching were corrected as previously described (68). Recovery curves were fitted using Graphpad Prism as previously described (68).

#### Acyl-Biotin Exchange (ABE)

ABE was performed according to published protocols, with modifications (39, 70). Lysates were prepared from 20-24 wandering L3 larvae per genotype by homogenizing the anterior half of the carcass after removing the gut and fat body, in lysis buffer (150mM NaCl, 50mM Tris, 5mM EDTA, pH 7.4) containing 1% Triton X-100 and protease inhibitors (Thermo). Following protein concentration measurement by BCA assay (Thermo), 200µg of protein per genotype was treated with 10mM TCEP, pH 7.4 (Thermo) and 4% SDS for 30 minutes at room temperature to reduce disulfide bonds and denature proteins. Samples were then treated with 30mM N-ethylmaleimide (NEM, Thermo) for 3 hours at room temperature, to cap newly exposed cysteines. Samples were then buffer exchanged with lysis buffer 4-5 times in 10K MWCO protein concentrator columns (Millipore) to remove residual NEM. Samples were then split 50:50 and half was treated with 0.8M hydroxylamine, pH 7.4 (Sigma) to cleave S-acyl groups for 1 hour at room temperature. The other half was diluted with an equivalent amount of lysis buffer and serves as a negative control. Samples were then buffer exchanged 3 times and treated with 1µM EZ-Link BMCC Biotin (Thermo) for 1 hour at room temperature. After buffer exchanging 3 times, biotinylated protein was purified using Pierce Streptavidin Magnetic Beads (Thermo) for 1 hour at room temperature or overnight at 4°C. Beads were washed twice and samples were eluted in 4X Laemmli sample buffer (Biorad) by boiling for 10 minutes. Biotin incorporation into proteins of interest was then analyzed by western blot. Western blotting was performed as described previously (71). Primary antibodies are listed in **Table 1**.

#### Image analysis and quantification

Image processing and quantification was performed using Fiji software (69). To quantify Baz particles, an approximately single-cell sized ROI of 104×104 pixels was defined. Baz particles in the ROI were then automatically segmented by creating binary masks from thresholded and smoothed images using the FeatureJ plugin. Segmented aggregates were then measured using the Analyze Particles function. To quantify aPKC levels, intensity was measured by drawing a line along the lateral membrane of a single cell medial section and using the measure function in Fiji. Mutant and WT cells were measured for each experiment and mutant cells were then normalized to the average intensity value of the corresponding WT data set. The resulting data were then analyzed using Microsoft Excel and Graphpad Prism 6. For significance in statistical tests: n.s.=p>0.05, *=p<0.05, **=p<0.01, ***=p<0.001 and ****=p<0.0001. Figures were assembled using Adobe Illustrator.

